# Effects of isochoric freezing on quality characteristics of raw bovine milk

**DOI:** 10.1101/2023.10.12.562068

**Authors:** Alan Maida, Cristina Bilbao-Sainz, Andrew Karman, Gary Takeoka, Matthew J. Powell-Palm, Boris Rubinsky

## Abstract

This study investigated the effects of isochoric freezing (IF) on the shelf-life and quality of raw bovine milk over a 5-week period. The results were compared with conventional refrigeration (RF) and refrigeration after pasteurization (HTST). The IF treatment process entailed storing liquid raw milk in isochoric chambers in thermodynamic equilibrium at -5°C /77MPa and -10°C /96MPa. Several parameters were analyzed, including microbiology count, physicochemical properties, indigenous enzyme activity, protein content, volatile organic compounds profile, and lipid degradation. Both raw and pasteurized milk experienced microbial level increases past the acceptable threshold (≥5.5 log CFU/mL) after 2 weeks and 5 weeks, respectively, leading to the deterioration of other parameters during storage. In comparison, microbiology count decreased significantly during storage for both IF treatment conditions but was more pronounced for the higher pressure (96MPa) treatment, leading to undetectable levels of microorganism after 5 weeks. IF treatment maintained stable pH, titratable acidity, viscosity, lipid oxidation, volatile profiles, total protein content, and lactoperoxidase activity throughout the storage period. Color was preserved during IF treatment at -5°C/77MPa; however, color was impacted during IF treatment at -10°C/96MPa. Protein structures were also modified during pressurized storage in both IF treatments. Overall, the study demonstrated that isochoric freezing could significantly increase the shelf-life of milk by reducing microbiology activity, whilst maintaining its nutritional content. These results underscore the potential role of isochoric freezing as a valuable tool in eliminating pathogens while maintaining quality characteristics similar to raw milk over long storage periods.

## Introduction

In light of a growing global population and the challenges posed by climate change and increasingly extreme weather patterns, ensuring food security has become a concern for the global community. Milk, renowned as one of nature’s most complete foods in terms of nutritional value [1], plays a pivotal role in addressing these insecurities. Concurrently, the alarming issue of food waste is taking a hefty toll. Across the globe, dairy farmers annually discard milk due to fluctuations in production, experiencing oversupply during peak seasons and shortages in off-peak periods. Other factors contributing to milk disposal include spoilage due to improper practices, a predicament experienced even in some of the most advanced dairy economies, where inappropriate handling lead to 29 million tons of dairy products being wasted due to fungal contamination and microbial attack annually [2]. The cumulative effect of these issues has led to a staggering global estimate of approximately 128 million tons of milk being discarded each year [3]. While current milk production meets consumer demand in many regions, the future presents uncertainties, driven by climate change and an increase in extreme weather patterns. Milk production is highly sensitive to heat stress, where intense summer heat has been shown to reduce milk production by up to 40% [4]. Considering the vital importance of milk and its by-products, such as baby formula, it is increasingly evident that improvements in preservation methods that may aid in waste reduction are needed for food security.

Milk, with its rich nutritional content and physicochemical properties, is an ideal breeding ground for bacteria, making milk one of the riskiest foods in terms of food safety [5]. Current preservation methods, primarily thermal treatment and freezing, offer partial solutions. High temperature-short time (HTST) pasteurization, for instance, successfully eliminates harmful microorganisms but extends milk’s shelf life only up to 14 days [6]. Another emerging method is ultra-high temperature (UHT) treatment, which is gaining popularity for its ability to sterilize milk by heating to 140°C for a couple of seconds [7]. Although this process extends the shelf life of milk to over 30 days, it comes at the cost of significant protein and amino acid modifications that can effect digestibility [8]. Freezing, while prolonging milk’s shelf life for months, significantly decreases the milk fat, protein and lactose contents [9]. Consequently, alternative methods are required to preserve milk.

Non-thermal processing technologies utilizing pressure have emerged as promising alternatives capable of reducing microbial load and thereby extending the shelf-life of milk. One such technique is high-pressure processing (HPP), which subjects milk to pressures ranging from 400 to 600 MPa for just a few seconds. HPP-treated milk, akin to conventionally pasteurized milk, requires refrigeration but exhibits the potential to prolong shelf-life for a substantial period, typically between 30 to 45 days [10]. Another innovative method, hyperbaric storage (HS), employs milder pressures, usually in the range of 20 to 150 MPa, for extended treatment durations. In a study by Duarte et al. [11], raw milk was treated at 75-100 MPa and room temperature for 60 days, resulting in microbial inactivation of endogenous microbial load, surrogate and pathogenic microorganisms, as well as bacterial spores. Remarkably, this approach also preserved the activity of heat-sensitive enzymes that exhibit antimicrobial activity [12]. However, under long-term hyperbaric storage, certain parameters influenced by temperature, such as volatiles and lipid oxidation, exhibited alterations [13]. These findings underscore the potential of mild pressures (75- 100 MPa) in both controlling microbial growth and preserving the biochemical activity of milk, but highlight the potential value of simultaneous optimization of storage temperature.

Isochoric freezing (IF) leverages confined aqueous thermodynamics to passively generate elevated pressures at mild sub-freezing temperatures, producing a technologically simple means of reducing metabolic processes while avoiding formation of ice in the stored food product [14,15]. The process involves filling a rigid container with an aqueous solution and a food product, subjecting it to sub-freezing temperatures, and allowing the expansion of ice to passively pressurize the interior of the system. Per Le Chatelier’s principle, the rising pressure hinders further freezing, inducing a pressurized two-phase liquid-ice equilibrium with a stable ice fraction at any given sub-freezing temperature. The system continues to freeze and increase pressure as the temperature of the system decreases, until it reaches the triple point of ice 1h, ice III, and the liquid. For pure water, this point occurs at approximately -21°C and 210MPa, with approximately 55% of the volume being converted to ice. This controlled freezing process allows food products to be placed within the liquid region of the chamber, preventing ice crystal formation and the according biophysical injury. Previous research has demonstrated the effectiveness of IF in reducing microbial activity and preserving physicochemical properties in various food products, without compromising nutritional content [16–18].

This study aims to contribute to the growing body of knowledge on the potential of isochoric freezing to preserve biological materials. We investigated herein the physicochemical and microbiological changes resulting from isochoric freezing of raw milk at temperatures of -5°C and -10°C and respective pressures of 77 and 96 MPa, for storage up to 5 weeks. We assessed key parameters, including pH, titratable acidity, color, viscosity, lactoperoxidase activity, free fatty acids, lipid oxidation, protein content, protein interactions, volatiles, and microbiology counts and compared them to raw milk and conventionally treated HTST milk. Our future research will delve into the effects of prolonged high-pressure exposure on milk shelf-life during subsequent refrigerated storage.

This work presents an initial exploration of the potential of isochoric milk preservation, offering a potentially promising solution to enhance food security, reduce food waste, and provide consumers with safe, nutritious milk after longer storage durations.

## 2. Materials and Methods

### 2.1. Isochoric system

Two different treatment conditions were used for isochoric experiments. Raw milk was treated at -5 °C/77 MPa and -10 °C/96 MPa using a pressure chamber made of Aluminum-7075 with a type-II anodize coating and a total volume capacity of 1500 mL, pressure-rated for up to 275MPa (BioChoric Inc., Bozeman, MT, USA). Each chamber was connected to a pressure gauge to monitor the pressure over time. The chambers were cooled using a chest freezer (Magic Chef Model #HMCF9W3, *MC Appliance Corporation,* Wood Dale, IL).

### 2.2. Experimental protocol

Raw bovine milk was purchased from a local market located in Albany, California. Four different conditions were compared: refrigerated raw milk (RF), refrigerated milk initially treated with high-temperature short-time (HTST) pasteurization, raw milk held under isochoric freezing (IF) at -5 °C/77 MPa, and raw milk held under IF at -10 °C/96 MPa. Initial tests and treatments began seven days before the labeled expiration date of the milk. Refrigerated (RF) raw bovine milk was used as the control. For the HTST treatment, raw bovine milk was heated to 72 °C and pasteurized for 15 seconds using a multipurpose sterilization unit (UHT/HTST Lab Microthermics model 25EHVH, Raleigh NC). The pasteurized milk was collected and divided aseptically into 3 sterilized polypropylene bottles through the drain out line. The initial quality of both RF and HTST milk was immediately tested to measure microbiological activity, physiochemical properties, and retention of certain compounds. The samples were then stored in a refrigerator (4 °C) for up to 5 weeks.

For the IF treatments, samples of the milk were prepared on the same day as the RF and HTST samples. Three bags of about 150 mL of raw bovine milk each were processed for each treatment. The sterilized bags were heat sealed with negligible air bubbles within and transferred to four separate isochoric chambers (total of twelve bags) filled with cold water. Cold water helped prevent accelerated bacteria growth as the chambers cooled to their preset temperatures in the chest freezers. After the treatment, the chamber was placed in a freezer set at 5 °C for 14h to melt all the ice within the chamber. Milk samples were treated under isochoric conditions for 2 and 5 weeks.

### 2.3. Microbiology

Decimal dilutions were prepared for total plate count and *Pseudomonas.* Appropriate dilutions were plated on plate count agar (PCA) for total aerobic mesophiles (TAM) and incubated at 30 °C for 3 days (ISO 4833:2013).

Numbers of *Pseudomonas* spp. were determined by spread plating appropriate dilutions on *Pseudomonas* agar base with CFC supplement (Oxoid Ltd., Basingstoke, Hampshire, U.K.) and incubated at 25 °C for 72h.

### 2.4. pH/Titration

The pH of the milk was determined in triplicate using a pH meter (Hanna instruments, USA).

The titratable acidity (TA) of the milk samples was determined by titrating 10 mL of diluted milk (4 mL of milk in 6 mL of DI water) to pH of 8.4 with 0.01 M sodium hydroxide solution. The results were expressed as grams of lactic acid per liter of milk based on:

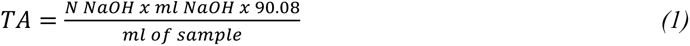

### 2.5. Color

The color of the bovine milk was measured using a tristimulus colorimeter (CM508D, Konica-Minolta Inc., Ramsey, NJ, USA) with a sample holder (CM-A128) and an 8 mm diameter target mask (CM-A195). The milk sample (10 mL) was pipetted into the sample holder covered with the black background and the color for each sample was measured three times. Results were expressed as L* (lightness), a* (redness/greenness), and b* (yellowness/ blueness) in the CIE Lab system. These values were used to calculate the overall color difference (ΔE*) according to the following equations:

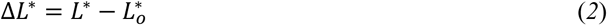

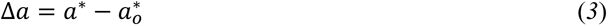

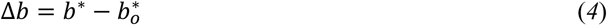

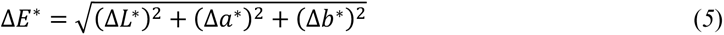

where L_o_*, a_o_*, and b_o_* represent the initial values prior to storage/treatment at day 0.

### 2.6. Turbidity

Turbidity of milk samples was monitored using a Hach Model TL2300 turbidimeter (Hach Company, Loveland, CO). The instrument was calibrated using StablCal® calibration standards (Hach Company), ranging between <0.1 and 4,000 nephelometric turbidity units (NTU). Diluted samples (1 mL of milk in 29 mL of water) were analyzed in a clean sample cell.

### 2.7. Viscosity

Milk viscosity measurements were performed on a DHR-3 rheometer (TA Instrument, New Castle, DE) at 20°C. A concentric cylinder geometry (28.04 mm bob diameter and 21.10 mm bob length) was used for testing. Three replicates were performed for each sample. For each replicate, 8 mL of milk was pipetted into the concentric cylinder cup and equilibrated at 20 °C for three minutes. The shear rate was increased logarithmically from 0.1 to 200 s^−1^. The viscosity of the sample was determined at a shear rate of 100 s^−1^.

### 2.8. Lipid Oxidation

Lipid oxidation was evaluated by quantification of secondary lipid oxidation products with the thiobarbituric acid reactive substances (TBARS) method, which was adapted from the method of Vyncke (1970). Briefly, 1 mL of milk was mixed with 1 mL of 7.5% trichloroacetic acid in water (TCA) in a 10 mL Falcon tube to allow the release of malondialdehyde (MDA) from the matrix and to precipitate some proteins. This was followed by centrifugation at 6000 rpm for 15 min at 4 °C. Subsequently, 1 mL of the supernatant was added to1 mL of 2-thiobarbituric acid (20 mM TBA in 99 % acetic acid glacial), slightly shaken and heated in a boiling water bath (100 °C for 40 min) and immediately cooled on ice for 5 min. The absorbance of the resulting pink reaction product was measured at 538 nm and room temperature (20 °C) (Microplate Spectrophotometer Multiskan GO, Thermo Scientific, Waltham, Massachusetts, USA) against a blank containing TCA instead of the sample. Measurements were performed in triplicate. For quantification, a standard curve of MDA was obtained from the hydrolysis of 1,1,3,3-tetramethoxypropane (Sigma Aldrich), ranging from 0.5 to 10.0 µM MDA. The results were expressed as µg MDA/mL of milk.

### 2.9. Protein Content

The total protein content was quantified according to the Lowry Method. The samples were diluted twice, (1:9 v/v) and (1:8 v/v), with 0.9% saline solution. The method was performed according to Waterborg & Matthews [19]. A calibration curve (R^2^=0.97) using a protein standard from 50 to 2000μg/mL was used to quantify the protein content.

### 2.10. Sodium dodecyl sulfate-polyacrylamide gel electrophoresis (SDS-PAGE)

The SDS-PAGE analyses were conducted with or without a reducing agent. For nonreduced sample preparation, 5 μL of milk was diluted with 50 μL of Tris-Glycine Native Sample Buffer (Novex, Thermo Fisher Scientific, Waltham, MA) and 45 μL of milli-Q water in a 2-mL plastic tube. For reduced samples, 5 μL of milk was diluted with 50 μL of Tris-Glycine SDS Sample Buffer (Novex, Thermo Fisher Scientific) and 35 μL of milli-Q water. Ten μL of sample reducing agent (NuPAGE, Thermo Fisher Scientific) was added to the mixture. Prepared reduced samples were shaken for 10 s and incubated at 85 °C for 2 min. Samples were analyzed using 10 to 20% precast tris-glycine protein mini gels (Invitrogen, Thermo Fisher Scientific) and Mini- Protein Electrophoresis Cells (Invitrogen, Thermo Fisher Scientific). Prestained Protein Ladder (NuPAGE, Thermo Fisher Scientific) was used to identify the detected proteins. Twenty microliters of each sample were added into the injection port and ran at 120 V for 90 min. The gels were stained with SimplyBlueTM (Thermo Fisher Scientific) and then destained with milli-Q water.

### 2.11. Lactoperoxidase Activity

The lactoperoxidase (LPO) activity assay was performed based on the method described by Marín et al. [20]. Briefly, milk was mixed with a solution of 0.65 mM ABTS (in 0.1 M sodium phosphate buffer, pH 6.0) and left for 30 min at 20 °C. Then, 0.1 mM hydrogen peroxide was added and mixed quickly to initiate the reaction, with the absorbance (Abs_412nm_) measured for 1 min. The enzymatic activity was calculated as the slope of Abs increment as a function of time and expressed as ΔAbs_412nm_ AU/min. All enzymatic assays were performed in triplicate for each storage condition.

The residual activity was calculated by using:

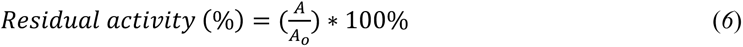

where *A* is the enzymatic activity in milk samples after storage and *A*_0_ represents the initial enzymatic activity prior to storage.

### 2.12. Volatile Organic Compounds

Volatile organic compounds (VOCs) of milk samples were measured using headspace solid-phase microextraction (HS-SPME) techniques. Sample preparation methods closely followed procedures described by Lemos et al. [13] with few modifications. Briefly, 10 mL of milk, 3.7g of sodium chloride, 250 µL of internal standard (aqueous cyclohexanone solution at 25 µg/mL), and a magnetic stir bar were added to a 240 mL screw top amber vial and sealed with a PTFE/silicone septum (22 mm × 2.54 mm thickness) and plastic screw cap combination. Sample vials were subsequently equilibrated in a water bath at 50 °C and gently stirred using a magnetic stir plate (80 rpm). Prior to extraction, the SPME fiber (DVB/CAR/PDMS, 50/30 µm coating, 2 cm length) was heated in the injection port (250 °C) of an Agilent 7890B gas chromatograph for 15 min.

After temperature equilibration of the sample vial, the SPME fiber was exposed to the headspace for 30 min while stirring. Fibers were then desorbed for 10 min (splitless) on a gas chromatograph (7890B) coupled to a mass selective detector (5977A MSD; Agilent Technologies, Palo Alto, USA). The injector temperature was set at 250 °C. A 60 m × 0.25 mm df = 0.25 µm DB-1 fused silica capillary column was used using helium as a carrier gas at constant flow of 1 mL/min (99.999% purity). The GC oven was programmed with the following conditions: 35 °C (5 min), ramped at 4°C/min until 100 °C, a second ramp at 10°C/min until 220 °C, and finally held at 220 °C (5 min). The purge valve to split vent was opened after 1 min at a flowrate of 30 mL/min following sample injection. The total run time lasted approximately 45 min. The MS transfer line and ion source were maintained at 250 °C and 230 °C, respectively. The MS quadrupole temperature was set to 150°C with an electron ionization of 70 eV, which was set in full scan mode (m/z 20 to 350 at 1.6 scan/s).

MassHunter Unknowns Analysis software (v. B.07.01) was used for data processing using the deconvolution algorithm. Identification of the VOCs was based on a combination of computer matching with mass spectral libraries (NIST 14) and retention index values derived from an n-alkane solution (c5 to n-c18) (Sigma Aldrich Co.). Only compounds with match factor scores ≥ 70 and signal-to-noise ratios ≥ 10 were selected for quantitation. Semi-quantitation was performed using cyclohexanone as the internal standard (IS). Results are expressed in µg eq. IS per 100 mL of milk.

### 2.13. Statistical Analysis

All experiments and analyses were carried out in triplicate. Analysis of variance was performed under all the different storage conditions, followed by a multiple comparison post hoc test, Tukey’s HSD test, at a 5% level of significance. Additionally, principal component analysis (PCA) was performed in order to identify the statistical patterns in VOC data.

## 3. RESULTS AND DISCUSSION

### 3.1. Microbiological evaluation

Given raw milk is not inherently sterile, the initial steps involved conducting microbiological assessments to examine how the different treatments and storage conditions affected the levels of indigenous microorganisms. For a general evaluation, the total bacteria count (TBC) and Pseudomonas spp. (PS) microbiological quality indexes were examined. Table 1 shows microbial populations found in RF, HTST, and IF-treated milk samples. Initial TPC and PS of raw milk were 6.98 ± 0.05 log CFU mL^-1^ and 7.22 ± 0.13 log CFU mL^-1^, respectively. After 2 weeks, the RF sample presented higher counts for both TPC and PS, 8.26 ± 0.19 and 8.07 ± 0.09 log CFU mL^-1^, respectively. Both initial and storage values were consistent with a previous study [21]. The RF samples in this study were well past its microbial shelf-life after 2 weeks (≥5.5 log CFU mL^-1^), and no further analyses were performed for these samples. The milk sample immediately after HTST treatment reached counts below 1 log CFU mL^-1^ (detection limit) for both TPC and PS, demonstrating HTST’s effectiveness in deactivating microorganisms. Interestingly, after 2 weeks of refrigerated storage at 4 °C, both TPC and PS for the HTST sample had increased by more than 5 log CFU mL^-1^. As expected, the HTST sample stored for 5 weeks at 4°C contained higher microbial load levels well above the acceptable threshold (≥5.5 log CFU mL^-1^). A previous study reported similar results for refrigerated HTST treated milk [6].

**Table 1:** Effects of microbial inactivation treatment and storage conditions on total aerobic mesophilic bacteria (TPC), and *Pseudomonas* spp. (PS) in raw bovine milk.

|  | Initial | RF | HTST |  |  | IF (-5°C) |  | IF (-10°C) |  |
| --- | --- | --- | --- | --- | --- | --- | --- | --- | --- |
|  | 0 Weeks | 2 Weeks | 0 Week | 2 Weeks | 5 Weeks | 2 Weeks | 5 Weeks | 2 Weeks | 5 Weeks |
| TPC | 6.98 | 8.26 | <0.01 | 5.28 | 9.59 | 3.60 | 1.33 | 1.26 | <0.01 |
| (Log CFU mL<sup>-1</sup>) | (.05) | (.19) |  | (.05) | (.06) | (.12) | (.28) | (.24) |  |
| PS | 7.22 | 8.07 | <0.01 | 5.31 | 9.80 | 4.85 | 1.80 | 3.45 | <0.01 |
| (Log CFU mL<sup>-1</sup>) | (.13) | (.09) |  | (.20) | (.13) | (.26) | (.040) | (.04) |  |

Results presented as mean and standard deviation in parenthesis.

Isochoric freezing affects microbial viability by potentially disrupting non-covalent bonds and cell membranes at high pressures. Prolonged exposure to these pressures can ultimately result in cell destruction and microbial deactivation [22]. After 2 weeks, the IF (-5 °C) treatment reduced TPC and PS counts to 3.60 ± 0.12 log CFU mL^-1^ and to 4.85 ± 0.26 log CFU mL^-1^, respectively. The IF(-10 °C) treatment had even greater reductions in TPC and PS counts after 2 weeks, 1.26 ± 0.24 and 3.45 ± 0.04 log CFU mL^-1^, respectively. After prolonged exposure to pressure, microbial counts decreased even further. After 5 weeks, IF(-5 °C) treatment resulted in more than 5 log CFU mL^-1^ reduction for both TPC and PS counts . For the IF(-10 °C) treatment after 5 weeks, both TPC and PS values were below the detection limit. The positive effect of isochoric freezing on the reduction of native microbial populations had been reported in a previous study [18]. A significantly higher reduction was achieved at higher pressures of IF(-10 °C) treatment compared to IF(-5 °C) treatment. Erkmen and Doğan [23] also found that HPP at 400 MPa and 600 MPa for 10 min reduced TPC in raw milk by 2 and 5 log CFU/ml, respectively. Another study showed that HPP treated homogenized milk stored at 4 °C had a microbiological shelf-life similar to that of pasteurized (90 °C for 15 s) milk [24]. This study demonstrated that within just two weeks, IF treatment significantly reduced microbial load levels below those of both RF (refrigeration) and HTST treated milk. These findings suggest that IF treatment could extend the microbiological shelf-life beyond that of untreated and pasteurized milk.

### 3.2. pH and titratable acidity

The pH and titratable acidity of the different milk samples under different storage conditions are shown in Table 2. Initial pH value of raw milk was 6.84 ± 0.04, which was not significantly different from the initial value of 6.82 ± 0.01 following HTST treatment. These values are similar to reported values in literature [25]. After two weeks of storage, the pH value of the RF sample decreased significantly to 6.57 ± 0.03, while the pH value of the HTST treated milk remained relatively stable at 6.80 ± 0.01. The decrease in pH value of the milk sample was probably caused by the accumulation of lactic acid produced by lactic acid bacteria. A previous study found that a decrease in pH value only occurred when microbial levels reached >6.5 log CFU mL^-1^ [25], which was consistent with the microbial numbers reported in Table 1. For IF treated samples, the pH values after 2 weeks at -5 °C and -10 °C showed no significant changes, with pH values of 6.77 ± 0.01 and 6.79 ± 0.02, respectively. Kim et al. [26] obtained comparable results with high pressure (200 MPa) and low temperature (-4 °C) treatments of raw milk for 10, 20, and 30 min. The authors found the treatments had no significant effects on the pH levels in the milk samples. After 5 weeks, the pH value in HTST treated milk had decreased significantly to 6.63 ± 0.0, which could be explained by the microbial count of >6.5 log CFU mL^-1^. For IF treatments, the pH values after 5 weeks at -5 °C (6.77 ± 0.01) and -10 °C (6.79 ± 0.02) showed no significant changes compared to the initial values. These results were consistent with the microbial count results, with both samples having microbial levels well below 6.5 log CFU mL^-1^.

**Table 2:** pH and titratable acidity (TA) values of raw bovine milk stored under different storage conditions.

|  | Initial | RF | HTST |  |  | IF (-5°C) |  | IF (-10°C) |  |
| --- | --- | --- | --- | --- | --- | --- | --- | --- | --- |
|  | 0 Weeks | 2 Weeks | 0 Week | 2 Weeks | 5 Weeks | 2 Weeks | 5 Weeks | 2 Weeks | 5 Weeks |
| pH | 6.84<br>(.04)<sup>a</sup> | 6.57<br>(.03)<sup>c</sup> | 6.82<br>(.01)<sup>a</sup> | 6.80<br>(.01)<sup>ab</sup> | 6.63<br>(.01)<sup>c</sup> | 6.77<br>(.01)<sup>b</sup> | 6.77<br>(.01)<sup>b</sup> | 6.79<br>(.02)<sup>ab</sup> | 6.79<br>(.02)<sup>ab</sup> |
| TA<br>(g LA/mL) | 1.27<br>(.08)<sup>c</sup> | 2.06<br>(.03)<sup>a</sup> | 1.36<br>(.08)<sup>c</sup> | 1.35<br>(.05)<sup>c</sup> | 1.69<br>(.02)<sup>b</sup> | 1.37<br>(.02)<sup>c</sup> | 1.58<br>(.06)<sup>b</sup> | 1.53<br>(.09)<sup>b</sup> | 1.63<br>(.06)<sup>b</sup> |

Results presented as mean and standard deviation in parenthesis. Different letters across each row indicate significant differences.

The initial titratability acidity (TA) value for raw milk was 1.27 ± 0.08 g/L lactic acid, which was similar to the initial value of 1.36 ± 0.08 for HTST treated milk. After 2 weeks, the TA value for the RF sample showed a significant increase to 2.06 ± 0.03 g/L, whereas the TA value for the HTST treated sample remained at 1.35 ± 0.05 g/L. These values were consistent with those found in literature [11]. Also, the TA values after 2 week IF treatment at -5°C (1.35 ± 0.05 g/L) and -10°C (1.37 ± 0.02) showed no significant changes compared with the initial value. After 5 weeks, the TA values for all milk samples increased significantly (*p*<0.05), albeit at different rates. The TA value for the HTST sample reached 1.69 ± 0.02 g/L, whereas the TA values for the IF(- 5°C) and IF(-10°C) samples reached 1.58 ± 0.06 g/L and 1.63 ± 0.06 g/L, respectively. The increase in TA values might be due to the continual growth of lactic acid bacteria (LAB), leading to more acidic conditions. A previous study reported a good correlation between acidity and microorganism growth [27]. For IF treatments, a possible explanation could be that due to the deactivation of LAB and the slight decrease in pH, the TA slightly increases.

### 3.3. Color and turbidity

Appearance is the first relevant attribute that determines whether a product is accepted or rejected by consumers. The appearance of the raw milk samples is shown in Fig. 1. Fresh raw milk showed a thick separation between the cream and the serum. After refrigeration for 2 and 5 weeks, the milk showed precipitation that resulted in fewer particles being present in the soluble phase. The HTST milk refrigerated for 5 weeks showed this same appearance. These changes may be related to the spoilage of milk by microorganisms that contributed to protein coagulation and fat degradation. The fresh raw milk and IF (-5 °C) sample showed no perceived differences in visual appearance. In comparison, the IF (-10 °C) sample showed formation of small and slightly visible aggregates.

**Fig. 1.**
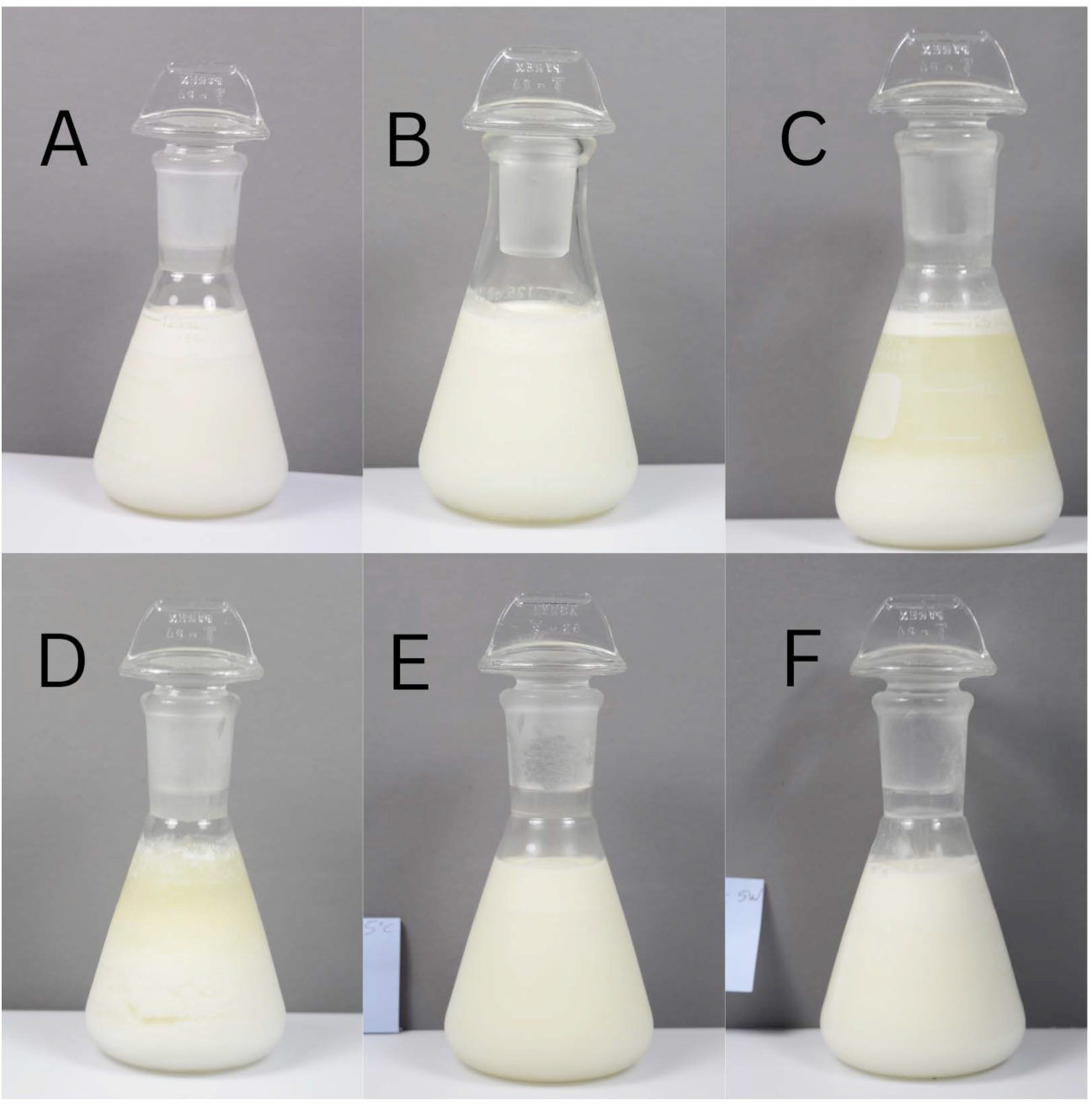
Photographs of (A) initial raw milk, (B) initial HTST treated milk, (C) RF after 5 weeks, (D) HTST after 5 weeks, (E) IF(-5°C) after 5 weeks, (F) IF(-10°C) after 5 weeks.

Table 3 shows the color parameters (L*, a*, b*) and turbidity values for all samples. Fresh raw milk had an L* value of 77.55 ± 0.15, an a* value of -1.78 ± 0.05, and a b* value of 3.38 ± 0.02. All samples undergoing different treatments and durations showed no significant changes (*p*<0.05) in L* values compared to the initial value. The natural color of milk (yellowish white) is ascribed to scattering and absorption of the visible light spectrum by dispersed fat globules, calcium caseinate, calcium phosphate, and pigments [28]. Previous studies found that L* tended to decrease significantly in value after HPP treatment [21]. This could be due to higher applied pressures in HPP treatment (300-600MPa) compared to IF treatment (75-100 MPa). Also, none of the treatments resulted in major changes in a* values (greenness). However, the b* (yellowness) value for the HTST sample significantly decreased to 0.03 ± 0.14 after 5 weeks. In comparison, the b* values for the IF (-5 °C) sample were comparable to that of the initial milk sample. Also, the b* values for the IF (-10 °C) sample at 2 and 5 weeks significantly increased in value to 4.86 ±.01 and 5.14 ±.15, respectively. These results were consistent with those reported by Stratakos et al. [21] and Lemos et al. [29] who also observed an increase in yellowness (increase of b∗) of raw milk treated with high pressures. They attributed the behavior to decreases in casein micelles and fat globule sizes due to high pressure that caused light to scatter [30].

**Table 3:**
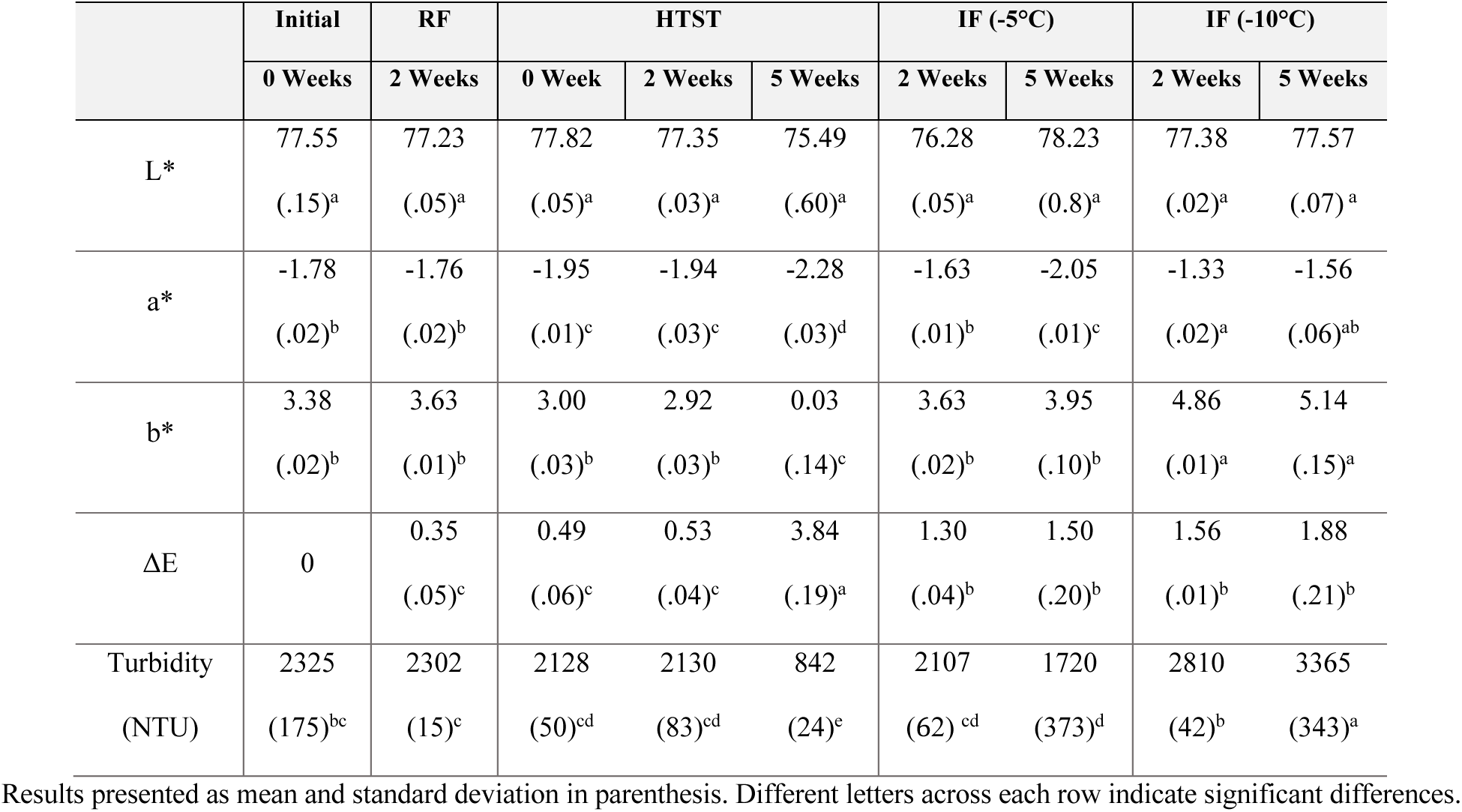
Chromatic parameters (L*, a*, b*, ΔE) and turbidity values of raw bovine milk stored using different storage methods.

The total color change (ΔE) values for the milk samples are shown in Table 3. ΔE is classified as not noticeable (0-0.5), slightly noticeable (0.5-1.5), noticeable (1.5-3.0), well visible (3.0-6.0), and great (6.0-12.0) [31] . According to this classification, the color difference of the HTST sample after 5 weeks is considered to be well visible to the human eye (3.84 ± 0.19). This was mainly caused by the increase in yellowness (b*) due to the increase in microbial loading (≥5.5 log CFU/mL). In comparison, the IF (-5 °C) sample had slightly noticeable color changes after 2 and 5 weeks, with values of 1.30 ± 0.04 and 1.50 ± 0.2, respectively On the other hand, the IF (-10 °C) sample had noticeable color changes after 2 and 5 weeks, with values of 1.56 ± 0.01 and 1.88 ± 0.21, respectively. The increase in total color change for the IF (-10 °C) sample at 5 weeks could be due to the formation of small aggregates.

The initial turbidity value for raw milk was 2325 ± 175 NTU (Table 3). HTST reduced milk turbidity due to alterations of the casein micelles [20]. The turbidity of the milk after IF (-5°C) treatment was significantly reduced to 2107 and 1720 NTU after 2 and 5 weeks, respectively. Previous authors also found a reduction in turbidity with pressure [32]. In contrast, the turbidity of the milk after IF (-10 °C) treatment increased to 2810 and 3365 NTU after 2 and 5 weeks, respectively. Milk turbidity was mainly related to the aggregation and dissociation of casein micelles [33]. The casein micelles are composed of casein molecules, colloidal calcium phosphate, and water molecules dispersed in the whey serum. Depending on the pressure and treatment time, casein micelles can either increase or decrease in size [34]. Disruption of casein micelles into submicelles under pressure could have reduced the turbidity of the IF (-5°C) milk samples. Needs et al. [35] suggested that the dissociation of casein micelles with pressure could be due to the disruption of hydrophobic and electrostatic interactions, and solubilization of colloidal calcium phosphate. The increase in turbidity levels in the IF (-10°C) sample might be due to an increase in casein micelle size from the formation of casein aggregates and complexation of denatured whey protein with casein micelles [34]. These changes in turbidity values were consistent with ΔE results, where the IF (-10 °C) sample had noticeable color change for both treatment time periods.

### 3.4. Viscosity

A comparison of the viscosity values at 100 s^-1^ between different treatments is shown in Fig. 2. The initial viscosity of fresh raw milk was 3.56 ± 0.01 mPa∗s. The HTST sample showed a slight decrease in milk viscosity to 3.38 ± 0.01 mPa∗s. The decrease in viscosity for heat treated samples had been attributed to a decrease in particle size when milk samples were heated to 60-70 °C [36]. The increase in viscosity values of HTST samples during storage might be due to microbial spoilage. IF (-5°C) treatment did not cause a clear effect on viscosity values. In comparison, IF (-10°C) treatment resulted in significant increases in viscosity, with the largest viscosity value of 3.71± 0.02 mPa∗s after 2 weeks. Previous studies showed increases in viscosity of milk with application of pressure [29,37,38]. The authors in the studies attributed the viscosity increase to the high pressures causing changes in the shape of casein micelles from roughly spherical to irregularly shaped particles due to significant disruption of micelles and denaturation of whey proteins [34].

**Fig. 2.**
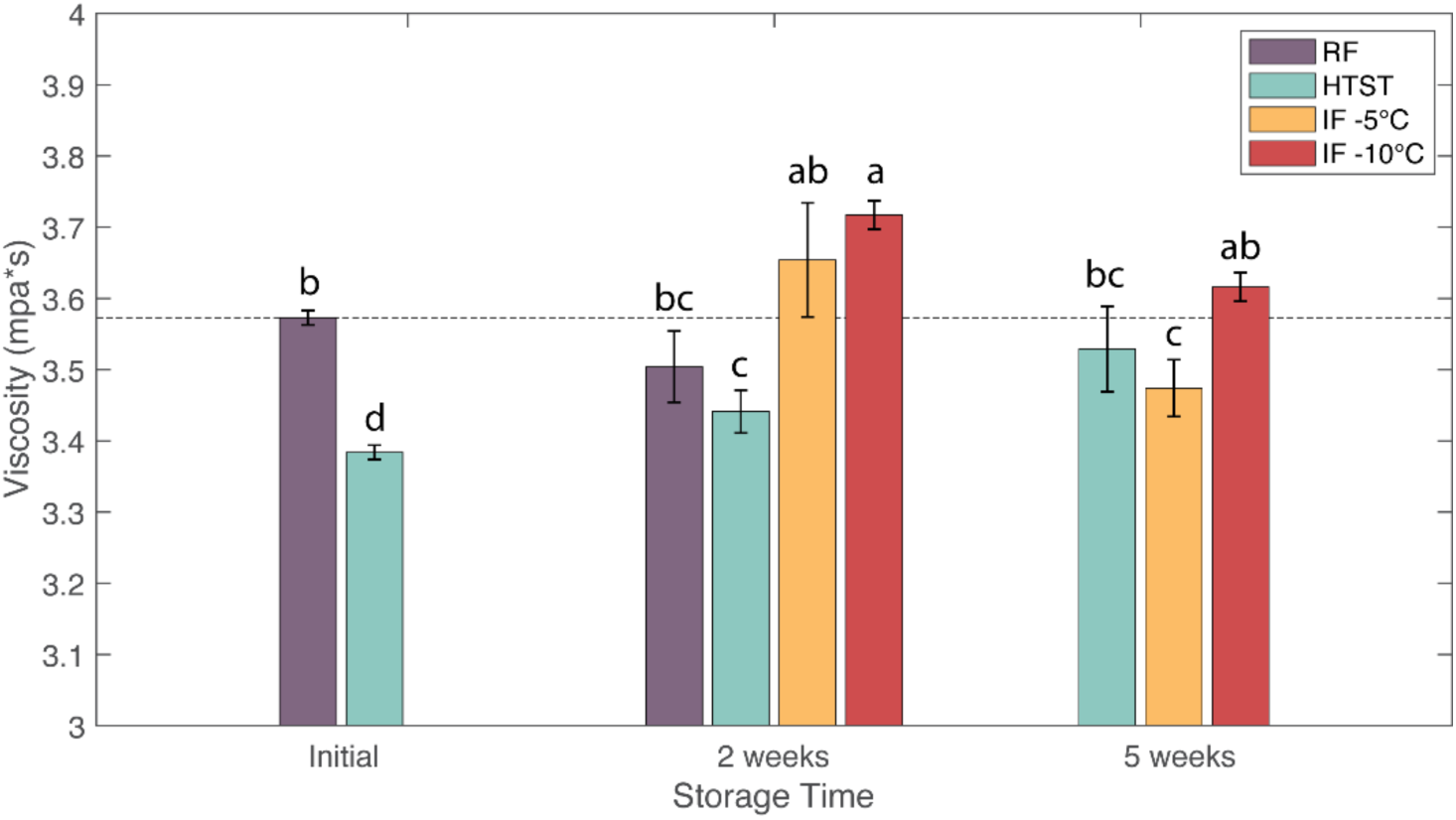
Effects of refrigeration (RF), HTST and isochoric freezing (IF) on viscosity values of raw bovine milk during storage. Values are the means of data sets and the standard deviations are indicated by the vertical error bars.

### 3.5. Lipid Oxidation

Lipid oxidation occurs in lipid-rich foods such as milk and is an indicator of oxidative rancidity. The initial TBARS value of raw milk was 0.62 ± 0.02 µg MDA/mL, consistent with values reported in literature [13]. Fig. 3 shows the TBARS values of the milk samples. Refrigeration caused about a 50% increase in lipid oxidation after 2 weeks. HTST samples also showed increased lipid oxidation during storage, reaching a value of 1.55 ± 0.19 µg MDA/mL (148% increase) after 5 weeks. This increase in lipid oxidation might be due to microbial and lipase activity. In comparison, IF treatments showed the lowest lipid oxidation values, with no significant differences compared to the fresh sample. After two weeks, TBARS values for IF(-5 °C) and IF(-10 °C) samples were 0.63 ± 0.16 and 0.86 ± 0.14 µg MDA/mL, respectively. After 5 weeks, these values remained relatively stable for the IF (-5°C) and IF (-10°C) samples at 0.69 ± 0.04 and 0.66 ± 0.04 µg MDA/mL, respectively. Previous studies on hyperbaric storage of milk at pressures of 75-100 MPa found higher overall TBARS values at room temperature (20 °C) storage for 60 days compared to refrigerated storage at 4 °C [13].

**Fig. 3.**
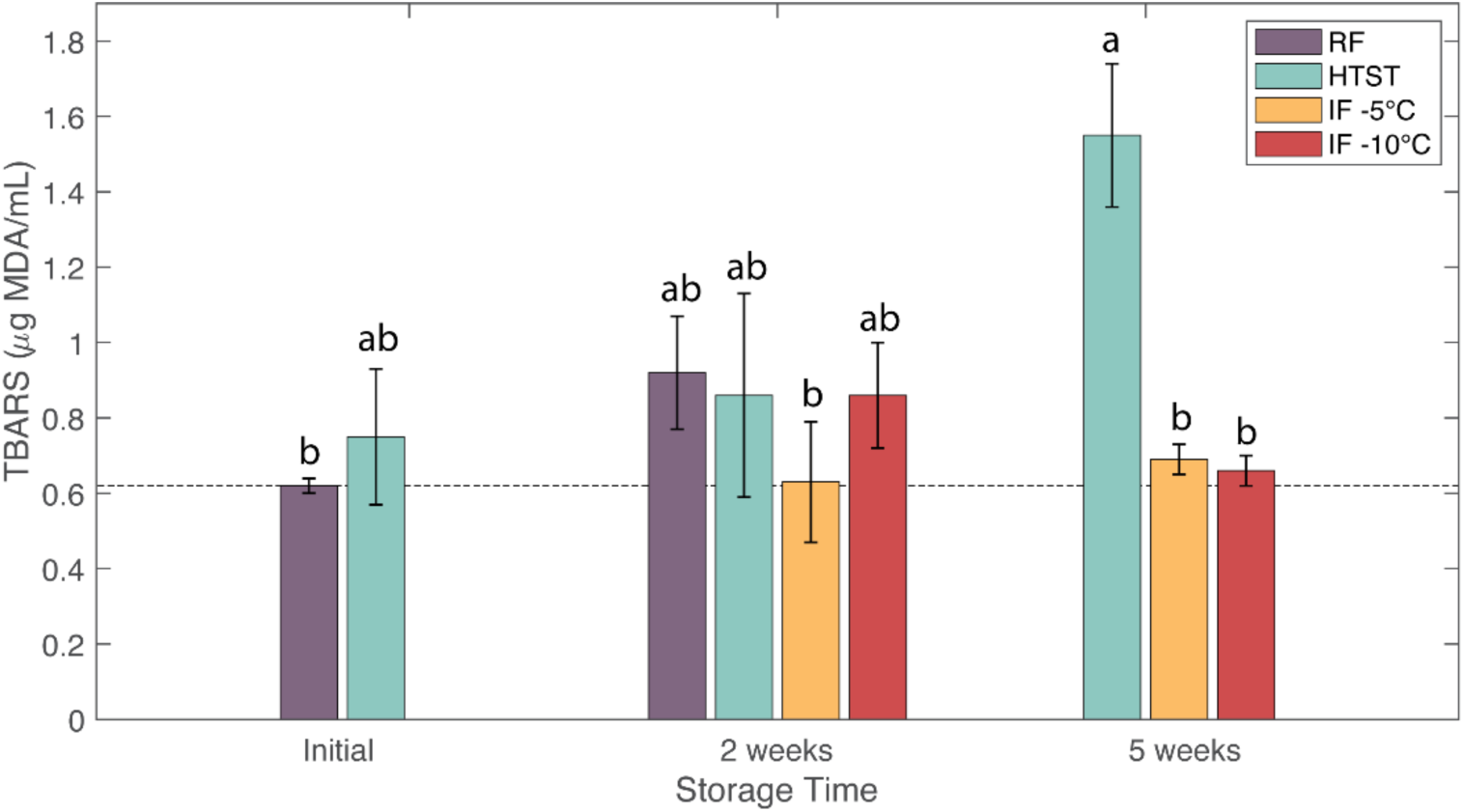
Effects of refrigeration (RF), HTST and isochoric freezing (IF) on lipid oxidation (μg MDA/mL) of raw bovine milk during storage. Values are the means of data sets and the standard deviations are indicated by the vertical error bars.

### 3.6. Protein Content and SDS-PAGE

The total protein content of all samples was determined by using the Lowry protein assay. The initial total protein content of raw milk was 32.16 ±3.33 mg/mL, similar to values reported in literature [39]. The total protein content in HTST treated milk showed an overall significant decrease in value to 23.53±2.33 mg/mL, which was attributed to an increase in proteolytic activity produced by the microflora of milk. In comparison, the protein contents after IF treatments showed no significant changes when compared with initial values and remained stable throughout the storage period. The total protein contents of the IF (-5 °C) sample were 30.57±.20 mg/mL after 2 weeks and 30.77±.42 mg/mL after 5 weeks. Also, the total protein contents of the IF (-10 °C) sample were 30.50 ±.66 mg/mL after 2 weeks and 32.17 ±.46 mg/mL after 5 weeks. A previous study also found that 60 days of hyperbaric storage at pressures of 75-100MPA did not affect the protein contents in milk [12].

Fig. 4 shows the SDS-PAGE results for the milk samples under native (A) and reducing (B) conditions. The protein bands for RF milk at week 2 and HTST treated milk at week 5 appeared with low intensity, indicating a decrease in soluble protein content attributed to an increase in the microbial load levels (Table 1). Surprisingly, under native conditions, all samples showed large molecular weight proteins near the top that could not diffuse into the gel as well as the absence of whey proteins. The α-casein and β-casein seemed to be unaffected by HTST treatment. Under reducing conditions, all the large molecular weight proteins disappeared, and the κ-casein and whey proteins appeared in the gels. This pattern was also previously observed in heat treated bovine milk [40]. The large aggregates that could not diffuse into the gels were composed predominantly of disulfide bonded α-la, β-lg, and κ-casein [41,42]. The IF (-5 °C) and IF (-10 °C) samples also displayed lower intensity for the casein bands under native conditions when compared to fresh milk. However, these bands became visible in the gels after the application of the reducing reagent. The generation of aggregates following high pressure treatments was previously reported by Jiang et al. [43]. β-lg is the most sensitive whey protein to pressure and can undergo denaturation under certain pressure [43,44]. Previous studies had found that pressure treatments between 50 and 100 MPa led to a predenatured state of β-lg, which can be reversible [45]. At higher pressures treatments, such as 400 MPa, can induce intermolecular-disulfide bonds between β-lg and *k*-casein. HPP treatment at longer durations or higher pressures can also induce interactions between β-lg and casein micelles that lead to sedimentation [34]. Under reducing conditions, the casein proteins appeared as sharp bands, suggesting that the denatured proteins did not sediment out of the solution. Huppertz et al. [34] reported that the level of sedimentable denatured β-lg increases with pressures greater than 100 MPa. under high pressure conditions (400-800 MPa) for 30 minutes, up to 80% of β-lg has been observed to sediment. The results of this study indicated that protein sedimentation remained limited in both IF (-5 °C) and IF (-10 °C), even after 5 weeks of storage.

**Fig. 4.**
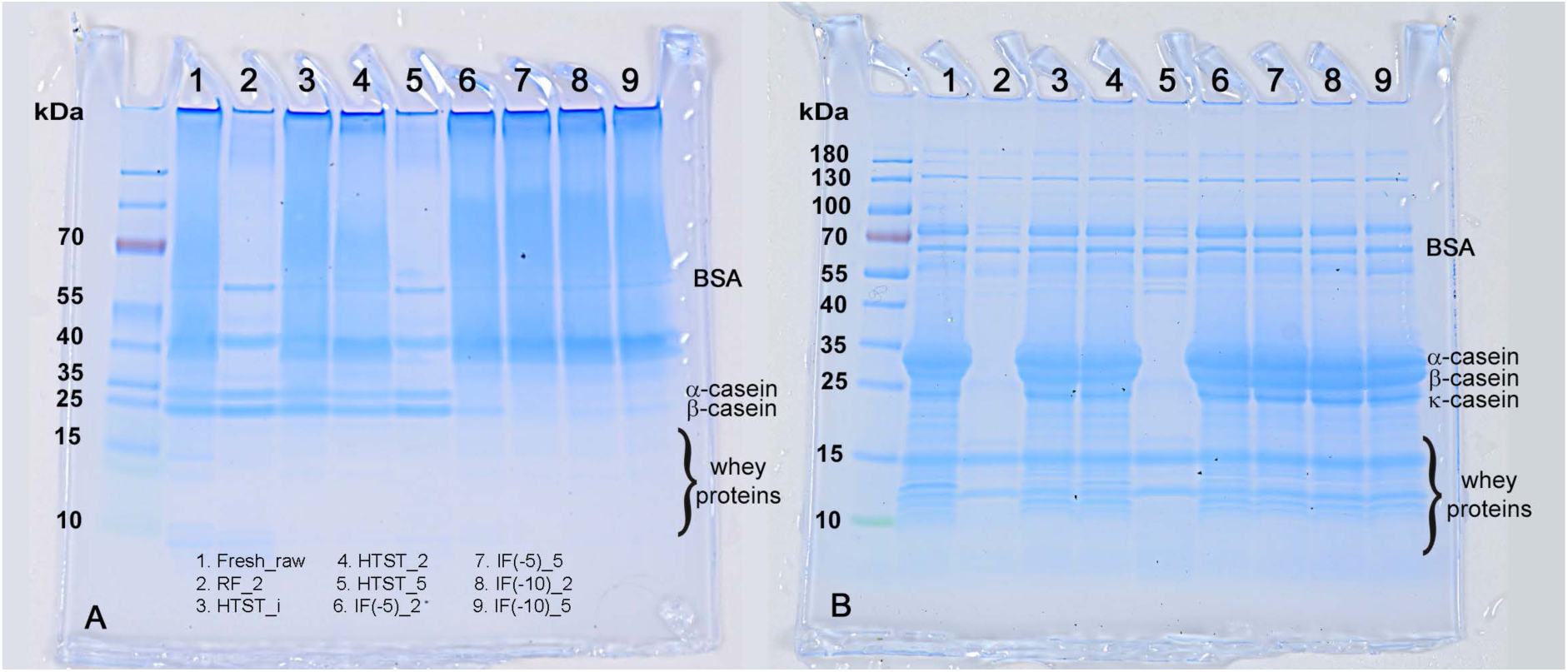
SDS-PAGE electrophoretic patterns of bovine milk casein after different treatments. Gel A (left) corresponds to SDS-PAGE patterns under native condition. Gel B (right) corresponds to reducing conditions.

### 3.7. Lactoperoxidase Activity

Lactoperoxidase (LPO) in milk catalyzes oxidation reactions in the presence of hydrogen peroxide to produce antimicrobial compounds. Enzyme activity is dependent on many variables such as pH, environment, pressure, and temperature [46]. All treated samples showed significant decreases in LPO activity when compared with fresh raw milk (Fig. 5), albeit to different degrees. HTST treatment immediately reduced LPO activity to 82% of the initial value. After 2 weeks, RF and HTST samples showed similar reductions in LPO activity to around 59% of the initial value. After 5 weeks, the HTST sample had a LPO activity that was 10% of the initial value. The IF (-5 °C) and IF (-10 °C) samples also showed significant decreases in LPO activity after two weeks to 72.5 and 80% of the initial value, respectively. After 5 weeks, the IF (-5 °C) sample showed no significant differences in LPO activity compared to 2 weeks, whereas the IF (-10 °C) sample showed a slight decrease to 64% of the initial value of fresh milk. Duarte et al. [12] also observed a decrease in activity when applying similar pressures at room temperature. More work is required, perhaps for longer treatment periods, to determine the specific effects of IF treatment on enzyme activity.

**Fig. 5.**
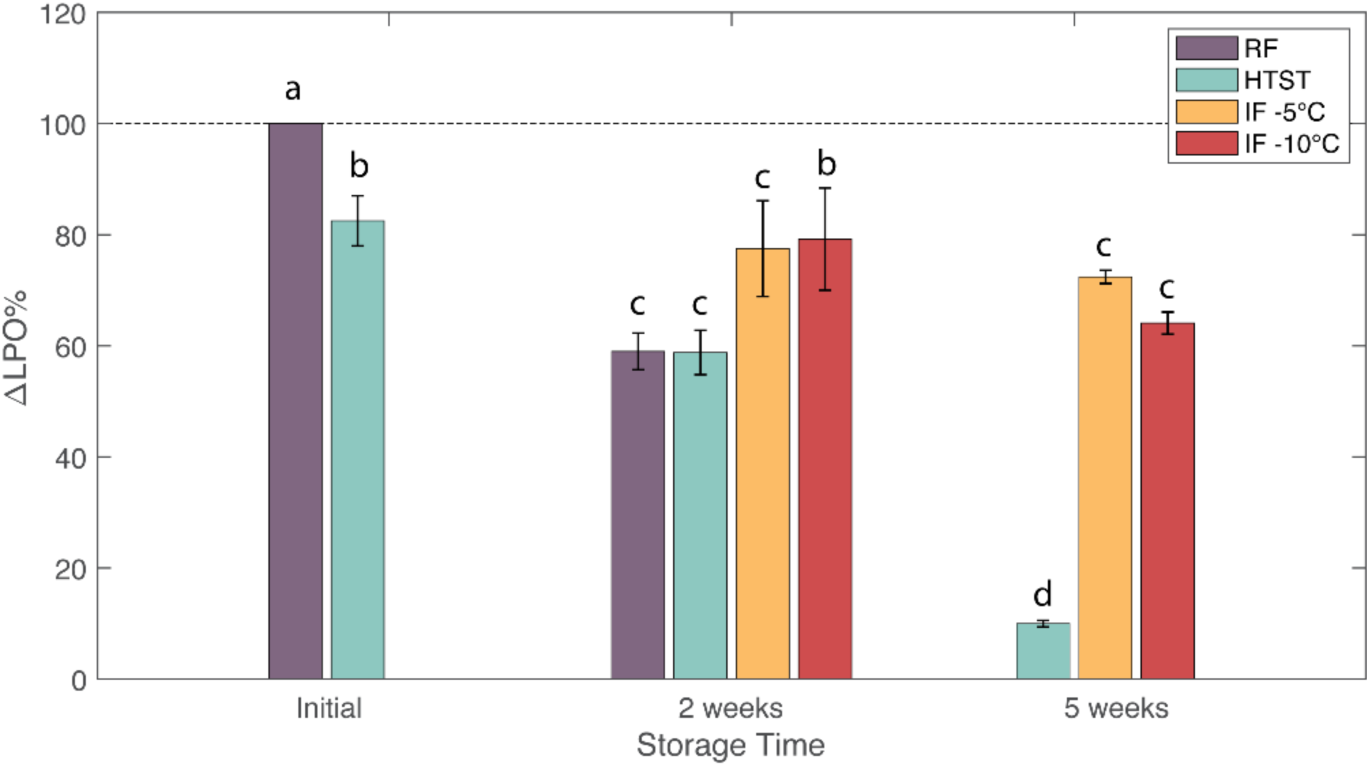
Effects of refrigeration (RF), HTST and isochoric freezing (IF) on lactoperoxidase activity of raw bovine milk during storage. Values are the means of data sets and the standard deviations are indicated by the vertical error bars.

### 3.8. SPME GC/MS Analysis

The volatile profile of milk samples of studied storage conditions was collected over a period of 5 weeks and the results are provided in Tables 4 and 5. In total, 33 compounds consisting of 5 acids, 5 alcohols, 7 aliphatic hydrocarbons, 2 aldehydes, 2 aromatics, 2 esters, 9 ketones, and 1 nitrogenous compound were identified. Of these, 28 compounds were confirmed through their corresponding reference literature retention indices while the remaining 5 compounds were tentatively identified using their mass spectra alone. Although the milk’s profile is believed to be partially influenced by diet (Wolf et al., 2013), the aroma composition found in the initial milk sample was consistent with literature findings [12,47].

**Table 4.** Volatile compounds (µg eq. IS/100 mL) grouped according to chemical class (acids, alcohols, aldehydes, aliphatic hydrocarbons, and aromatics) of raw milk stored up to 5 weeks. Storage conditions include refrigeration at 4°C (RF), pasteurization and subsequent refrigeration at 4°C (HTST), and isochoric freezing (IF) at –5°C and –10°C. Results presented as mean and standard deviation in parenthesis. Different letters across each row indicate significant differences.

| Volatile Compound | $RI^{ref}$ | $RI$ | Initial | RF | HTST | | | IF ($-5^{\circ}\text{C}$) | | IF ($-10^{\circ}\text{C}$) | |
| --- | --- | --- | --- | --- | --- | --- | --- | --- | --- | --- | --- |
|  |  |  | 0 Weeks | 2 Weeks | 0 Weeks | 2 Weeks | 5 Weeks | 2 Weeks | 5 Weeks | 2 Weeks | 5 Weeks |
| **Acids** |  |  | 2.87<br>(0.76) <sup>a</sup> | 32.53<br>(7.4) <sup>b</sup> | 4.66<br>(1.28) <sup>a</sup> | 2.55<br>(1.73) <sup>a</sup> | 149.15<br>(10.49) <sup>c</sup> | 3.6<br>(1.65) <sup>a</sup> | 3.67<br>(0.7) <sup>a</sup> | 1.93<br>(0.84) <sup>a</sup> | 1.63<br>(1.74) <sup>a</sup> |
| Butanoic acid | 795 | 785 | ND | 4.17<br>(1.52) | ND | ND | ND | ND | ND | ND | ND |
| Hexanoic acid | 973 | 971 | 2.33<br>(0.26) <sup>a</sup> | 16.94<br>(3.79) <sup>b</sup> | 2.61<br>(0.62) <sup>a</sup> | 1.38<br>(1.42) <sup>a</sup> | 51.33<br>(7.97) <sup>c</sup> | 2.49<br>(1.03) <sup>a</sup> | 2.46<br>(0.29) <sup>a</sup> | 1.53<br>(0.56) <sup>a</sup> | 0.98<br>(1.47) <sup>a</sup> |
| Octanoic acid | 1165 | 1156 | 0.54<br>(0.63) <sup>a</sup> | 8.97<br>(1.72) <sup>b</sup> | 1.76<br>(0.54) <sup>a</sup> | 1.18<br>(0.31) <sup>a</sup> | 55.04<br>(8.36) <sup>c</sup> | 1.11<br>(0.64) <sup>a</sup> | 1.21<br>(0.44) <sup>a</sup> | 0.40<br>(0.49) <sup>a</sup> | 0.65<br>(0.32) <sup>a</sup> |
| Decanoic acid | 1335 | 1344 | ND | 2.45<br>(0.44) <sup>a</sup> | 0.29<br>(0.12) <sup>a</sup> | ND | 41.46<br>(6.38) <sup>b</sup> | ND | ND | ND | ND |
| Dodecanoic acid | 1547 | 1539 | ND | ND | ND | ND | 1.33<br>(0.33) | ND | ND | ND | ND |
| **Alcohols** |  |  | 3.34<br>(0.57) <sup>b</sup> | 4.18<br>(0.62) <sup>b</sup> | ND | 1.39<br>(0.55) <sup>a</sup> | 1.21<br>(0.38) <sup>a</sup> | 1.21<br>(0.17) <sup>a</sup> | 0.91<br>(0.28) <sup>a</sup> | 1.32<br>(0.41) <sup>a</sup> | 0.73<br>(0.34) <sup>a</sup> |
| Ethanol | 440 | 466 | ND | 1.87<br>(0.38) | ND | ND | ND | ND | ND | ND | ND |
| 2-propanol | 475 | 493 | ND | 0.89<br>(0.06) | ND | ND | ND | ND | ND | ND | ND |
| 2-methyl-2-propanol | 512 | 510 | 2.40<br>(0.36) <sup>b</sup> | 1.16<br>(0.37) <sup>a</sup> | ND | 1.19<br>(0.54) <sup>a</sup> | 1.21<br>(0.38) <sup>a</sup> | 0.85<br>(0.15) <sup>a</sup> | 0.84<br>(0.34) <sup>a</sup> | 1.13<br>(0.37) <sup>a</sup> | 0.60<br>(0.31) <sup>a</sup> |
| 2-methyl-1-propanol | 607 | 613 | 0.65<br>(0.22) <sup>b</sup> | ND | ND | ND | ND | 0.13<br>(0.15) <sup>a</sup> | ND | ND | ND |
| 2-methyl-2-butanol | 614 | 622 | 0.30<br>(0.02) <sup>c</sup> | 0.25<br>(0.06) <sup>bc</sup> | ND | 0.19<br>(0.03) <sup>abc</sup> | ND | 0.23<br>(0.02) <sup>bc</sup> | 0.08<br>(0.09) <sup>a</sup> | 0.19<br>(0.06) <sup>abc</sup> | 0.13<br>(0.03) <sup>ab</sup> |
| **Aldehydes** |  |  | ND | ND | 1.51<br>(2.23) <sup>a</sup> | ND | ND | 0.65<br>(0.77) <sup>a</sup> | 0.81<br>(0.12) <sup>a</sup> | ND | 0.54<br>(0.62) <sup>a</sup> |
| Hexanal | 783 | 775 | ND | ND | ND | ND | ND | 0.65<br>(0.77) <sup>a</sup> | 0.81<br>(0.12) <sup>a</sup> | ND | 0.54<br>(0.62) <sup>a</sup> |
| Nonanal | 1082 | 1084 | ND | ND | 1.51<br>(2.23) | ND | ND | ND | ND | ND | ND |
| **Aliphatic Hydrocarbons** |  |  | 2.19<br>(1.53) <sup>a</sup> | 10.02<br>(2.78) <sup>cd</sup> | ND | 12.12<br>(3.8) <sup>d</sup> | 12.76<br>(1.71) <sup>cd</sup> | 8.94<br>(4.42) <sup>bcd</sup> | 4.91<br>(3.05) <sup>ab</sup> | 7.47<br>(1.12) <sup>bc</sup> | 7.16<br>(3.21) <sup>bc</sup> |
| Pentane | 500 | 500 | ND | 0.10<br>(0.11) | ND | ND | ND | ND | ND | ND | ND |
| Heptane | 700 | 700 | ND | ND | ND | ND | 0.15<br>(0.03) | ND | ND | ND | ND |
| 4-methyl-heptane | 768 | 768 | 0.82<br>(0.60) <sup>a</sup> | 3.10<br>(0.88) <sup>ab</sup> | ND | 3.94<br>(1.06) <sup>b</sup> | 4.02<br>(0.56) <sup>b</sup> | 2.75<br>(1.39) <sup>ab</sup> | 1.26<br>(1.58) <sup>ab</sup> | 2.47<br>(0.32) <sup>ab</sup> | 2.52<br>(1.05) <sup>ab</sup> |
| Octane | 800 | 800 | ND | 0.22<br>(0.26) <sup>a</sup> | 0.21<br>(0.05) <sup>a</sup> | ND | 0.40<br>(0.06) <sup>a</sup> | 0.41<br>(0.07) <sup>a</sup> | 0.36<br>(0.15) <sup>a</sup> | ND | ND |
| 2,4-dimethyl-heptane | 824 | 824 | ND | 0.17<br>(0.20) <sup>a</sup> | ND | 0.38<br>(0.19) <sup>a</sup> | 0.47<br>(0.08) <sup>a</sup> | 0.18<br>(0.21) <sup>a</sup> | ND | ND | ND |
| 2,4-dimethyl-1-Heptene | — | 840 | 1.38<br>(0.92) <sup>a</sup> | 6.07<br>(1.52) <sup>b</sup> | ND | 7.79<br>(2.58) <sup>b</sup> | 7.23<br>(1.08) <sup>b</sup> | 6.00<br>(2.91) <sup>b</sup> | 3.64<br>(1.70) <sup>ab</sup> | 4.89<br>(0.79) <sup>ab</sup> | 4.64<br>(2.16) <sup>ab</sup> |
| Decane | 1000 | 1000 | ND | 0.58<br>(0.22) <sup>b</sup> | ND | ND | 0.88<br>(0.07) <sup>b</sup> | ND | ND | 0.11<br>(0.13) <sup>a</sup> | ND |
| **Aromatics** |  |  | 0.44<br>(0.24) <sup>ab</sup> | 0.9<br>(0.17) <sup>bc</sup> | 0.12<br>(0.05) <sup>a</sup> | 1.09<br>(0.36) <sup>c</sup> | 2.48<br>(0.19) <sup>d</sup> | 0.94<br>(0.48) <sup>bc</sup> | 0.67<br>(0.08) <sup>b</sup> | 0.63<br>(0.19) <sup>bc</sup> | 0.76<br>(0.27) <sup>bc</sup> |
| Toluene | 753 | 751 | 0.29<br>(0.06) <sup>ab</sup> | 0.40<br>(0.05) <sup>b</sup> | 0.12<br>(0.05) <sup>a</sup> | 0.44<br>(0.09) <sup>b</sup> | 0.52<br>(0.07) <sup>b</sup> | 0.53<br>(0.23) <sup>b</sup> | 0.37<br>(0.04) <sup>ab</sup> | 0.32<br>(0.06) <sup>ab</sup> | 0.37<br>(0.08) <sup>ab</sup> |
| Benzene, 1,3-bis(1,1-dimethylethyl)- | — | 1250 | 0.16<br>(0.19) <sup>a</sup> | 0.50<br>(0.13) <sup>ab</sup> | ND | 0.66<br>(0.27) <sup>b</sup> | 1.96<br>(0.13) <sup>c</sup> | 0.41<br>(0.25) <sup>ab</sup> | 0.30<br>(0.09) <sup>ab</sup> | 0.31<br>(0.13) <sup>ab</sup> | 0.38<br>(0.19) <sup>ab</sup> |

**Table 5.** Volatile compounds (µg eq. IS/100 mL) grouped according to chemical class (esters, ketones, and nitrogenous compounds) of raw milk stored up to 5 weeks. Storage conditions include refrigeration at 4°C (RF), pasteurization and subsequent refrigeration at 4°C (HTST), and isochoric freezing (IF) at –5°C and –10°C. Results presented as mean and standard deviation in parenthesis. Different letters across each row indicate significant differences.

| Volatile Compound | $RI^{ref}$ | $RI$ | Initial | RF | HTST | | | IF (-5°C) | | IF (-10°C) | |
| --- | --- | --- | --- | --- | --- | --- | --- | --- | --- | --- | --- |
|  |  |  | 0 Weeks | 2 Weeks | 0 Week | 2 Weeks | 5 Weeks | 2 Weeks | 5 Weeks | 2 Weeks | 5 Weeks |
| **Esters** |  |  | ND | 1.59<br>(0.35) | ND | ND | ND | ND | ND | ND | ND |
| Ethyl acetate | 601 | 601 | ND | 1.33<br>(0.07) | ND | ND | ND | ND | ND | ND | ND |
| Hexanoic acid, ethyl ester | 986 | 983 | ND | 0.26<br>(0.30) | ND | ND | ND | ND | ND | ND | ND |
| **Ketones** |  |  | 12.37<br>(1.21) <sup>c</sup> | 8.64<br>(1.16) <sup>ab</sup> | 11.52<br>(1.01) <sup>bc</sup> | 7.58<br>(2.88) <sup>a</sup> | 11.83<br>(1.44) <sup>c</sup> | 11.16<br>(1.1) <sup>bc</sup> | 10.94<br>(0.23) <sup>abc</sup> | 11.47<br>(0.7) <sup>bc</sup> | 10.33<br>(2.05) <sup>abc</sup> |
| Acetone | 468 | 486 | 7.59<br>(0.98) <sup>d</sup> | 3.44<br>(0.75) <sup>ab</sup> | 6.64<br>(1.11) <sup>cd</sup> | 4.28<br>(2.00) <sup>abc</sup> | 2.44<br>(0.41) <sup>a</sup> | 6.97<br>(0.58) <sup>cd</sup> | 6.45<br>(0.19) <sup>cd</sup> | 7.33<br>(0.73) <sup>d</sup> | 6.05<br>(1.75) <sup>bcd</sup> |
| 2-butanone | 575 | 570 | 4.32<br>(0.50) <sup>c</sup> | 2.15<br>(0.17) <sup>a</sup> | 4.71<br>(0.17) <sup>c</sup> | 2.61<br>(0.94) <sup>ab</sup> | 1.24<br>(0.09) <sup>a</sup> | 4.19<br>(0.63) <sup>c</sup> | 3.94<br>(0.12) <sup>c</sup> | 3.92<br>(0.58) <sup>c</sup> | 3.73<br>(0.36) <sup>bc</sup> |
| 2-pentanone | 667 | 661 | ND | 0.59<br>(0.68) <sup>a</sup> | ND | ND | 1.12<br>(0.02) <sup>a</sup> | ND | ND | ND | ND |
| 3-methyl-2-butanone | 659 | 662 | 0.46<br>(0.06) <sup>a</sup> | 0.52<br>(0.60) <sup>a</sup> | ND | ND | ND | ND | 0.54<br>(0.14) <sup>a</sup> | 0.21<br>(0.25) <sup>a</sup> | 0.55<br>(0.11) <sup>a</sup> |
| 2,4-dimethyl-3-heptanone | — | 701 | ND | 0.06<br>(0.06) | ND | ND | ND | ND | ND | ND | ND |
| 2-Heptanone | 847 | 868 | ND | 1.10<br>(0.16) <sup>b</sup> | 0.17<br>(0.03) <sup>a</sup> | 0.29<br>(0.03) <sup>a</sup> | 3.18<br>(0.64) <sup>c</sup> | ND | ND | ND | ND |
| 4-methyl-2-heptanone | — | 920 | ND | 0.17<br>(0.20) <sup>a</sup> | ND | 0.40<br>(0.18) <sup>a</sup> | 0.41<br>(0.04) <sup>a</sup> | ND | ND | ND | ND |
| 2-Nonanone | 1094 | 1073 | ND | 0.61<br>(0.10) <sup>a</sup> | ND | ND | 2.54<br>(0.49) <sup>b</sup> | ND | ND | ND | ND |
| 2-Undecanone | 1279 | 1276 | ND |  | ND | ND | 0.88<br>(0.15) | ND | ND | ND | ND |
| **Nitrogenous Compounds** |  |  | 11.81<br>(2.32) <sup>ab</sup> | 15.32<br>(2.26) <sup>c</sup> | 9.71<br>(1.07) <sup>ab</sup> | 8.27<br>(0.66) <sup>ab</sup> | 14.89<br>(2.97) <sup>c</sup> | 10.34<br>(1.34) <sup>b</sup> | 7.91<br>(0.9) <sup>a</sup> | 10.01<br>(2.51) <sup>ab</sup> | 8.82<br>(0.49) <sup>ab</sup> |
| Oxime-, methoxy-phenyl_ | — | 890 | 11.81<br>(2.32) <sup>ab</sup> | 15.32<br>(2.26) <sup>c</sup> | 9.71<br>(1.07) <sup>ab</sup> | 8.27<br>(0.66) <sup>ab</sup> | 14.89<br>(2.97) <sup>c</sup> | 10.34<br>(1.34) <sup>b</sup> | 7.91<br>(0.90) <sup>a</sup> | 10.01<br>(2.51) <sup>ab</sup> | 8.82<br>(0.49) <sup>ab</sup> |

$RI^{ref}$ are literature retention indices obtained on a DB-1 column as reported by http://webbook.nist.gov
ND = not detected

The three most abundant compounds present in the initial milk sample was methoxyphenyl-oxime (36%), acetone (23%), and 2-butanone (13%). While there is little information regarding flavor characteristics of methoxyphenyl-oxime, its presence in milk has been reported elsewhere [13,47]. Acetone and 2-butanone, both ketones, are common milk aromas which might originate from the cow’s metabolism. Ketone aromas contribute to floral or fruity aromas in milk. The remaining compounds are classified as free fatty acids (FFAs), branched hydrocarbons, alcohols, and aromatics. In the context of flavor, FFAs are of particular importance. FFAs can result from various sources, such as in the lipolysis of milk triglycerides, lipid oxidation reactions, or microbial growth. Thus, high concentrations of (volatile) FFAs suggests degradation of milk quality and flavor.

After two weeks of storage, a significant increase (*p*<0.05) in FFAs (∼11×) was observed in the refrigerated samples (RF). In addition to the increase in all formerly identified FFAs, newly formed FFAs included butanoic acid and decanoic acid. Ethanol was also detected in the refrigerated raw milk samples only. The newly formed ester, ethyl acetate, contributes to the sourness of the milk and is an indicator of milk quality degradation. Significant increases in aliphatic hydrocarbons were also observed, however their impact to the milk’s flavor is considered secondary due to their high odor thresholds.

Same-day pasteurization of the initial milk sample did not result in substantial changes in nearly all chemical categories. However, small amounts of newly formed decanoic acid were detected. The formation of nonanal, which delivers sweet grass-like odors was also observed. Milk pasteurization followed by 2 weeks of refrigeration (HTST) led to a significant increase in aliphatic hydrocarbon content but a decline in overall ketone amounts. The net outcome of these changes likely signifies the onset of milk flavor degradation. However, if compared with RF results, HTST substantially improves preservation of milk flavor. By week 5, changes in HTST’s volatile profile were apparent as there was∼50-fold increase in free fatty acid content suggesting milk spoilage.

IF treatments resulted in an overall better preservation of raw milk VOC profiles, even after 5 weeks, with no notable changes (*p* > 0.05) observed for FFAs and esters when compared to their initial values. Milk aroma profiles were also measured following isochoric freezing at – 5°C and –10°C. Like the HTST findings, two weeks of IF storage resulted in relatively few significant changes in the aroma profile. The most significant change appears to be an increase in aliphatic hydrocarbons. Interestingly, the resulting aroma profiles between IF between –5°C and –10°C were highly similar. Remarkably, 5 weeks of IF storage led to no substantive changes in the milk’s volatile profile from their 2 weeks measurements. Such findings suggest IF effectively preserves milk flavor for time periods much longer than conventional pasteurization techniques.

A principal components analysis was performed on the VOCs in order to identify possible relationships between storage conditions. Fig. 6 plots the first two principal components (PC1 and PC2). The biplot shows that, apart from ketones, most chemical classes had high loadings in the first principal component. Summing the first two PCs results in 71.5% variance explained. As expected, isochoric samples are closely grouped to the initial milk sample.

**Fig. 6.**
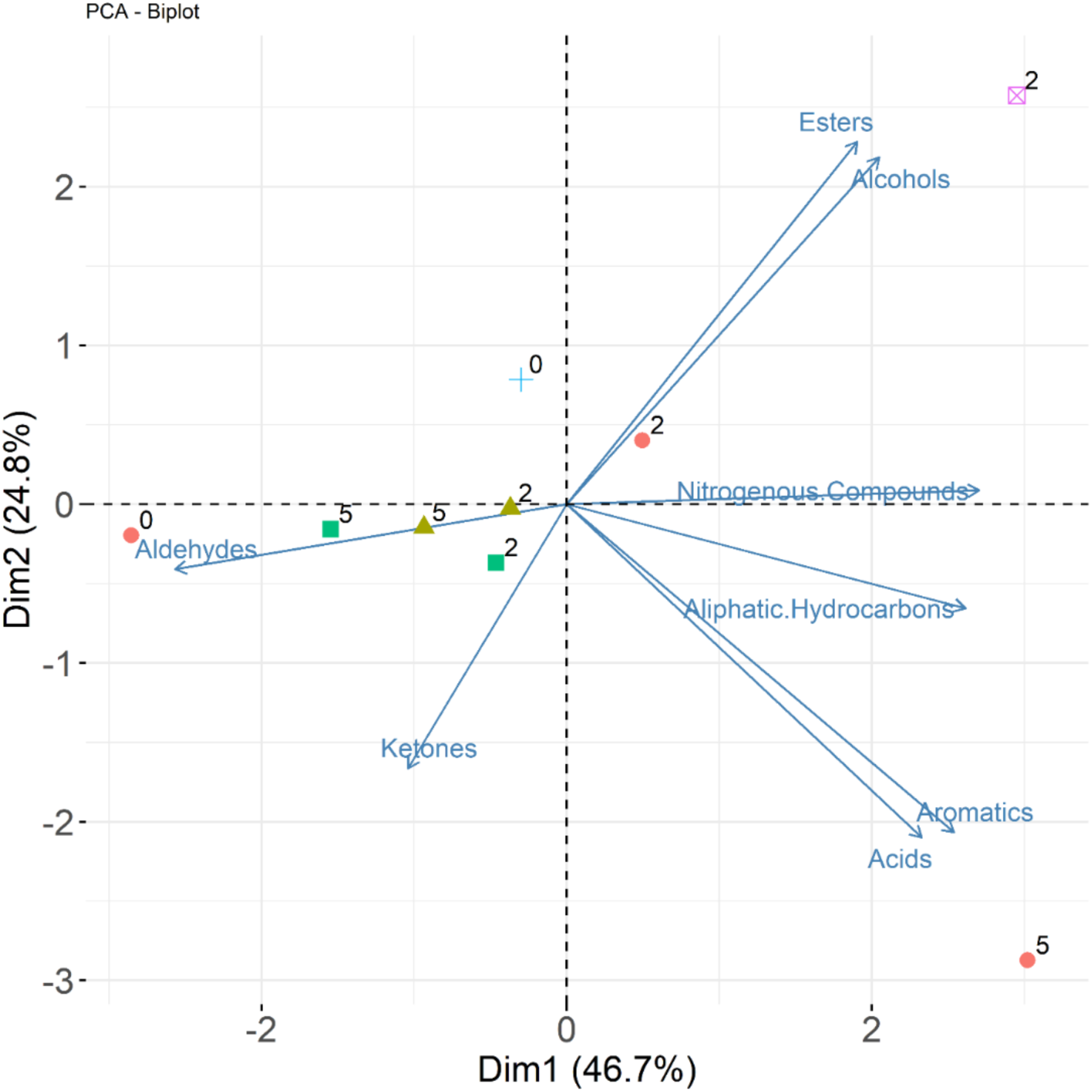
Principal component analysis of volatile samples in milk at initial conditions (+), isochoric freezing at –5°C (**▪),** isochoric freezing at –10°C (▴), pasteurization and subsequent refrigeration (●), and refrigeration only (⊠). Numbers adjacent to symbols correspond to number of weeks storage at each condition.

## Conclusions

This work describes the effects of isochoric freezing at -5°C/75MPa and -10°C/100MPa on the physicochemical properties, indigenous enzyme activity, protein content, volatile organic compounds profile, lipid degradation, and microbiological activity of raw bovine milk. Overall, prolonged storage under isochoric freezing conditions, when compared to refrigeration (RF), resulted in superior quality across measured parameters. Prolonged isochoric freezing was also able to better preserve milk for longer storage periods as compared to HTST treated milk. Unlike conventional refrigeration and HTST treatment, IF treated milk at -5°C/75MPa and -10°C/100MPa not only inhibited microbial growth, but also significantly reduced microbial count, with IF(-10°C) producing counts below the detectable limit. This anti-microbial effect, driven by the combination of enhanced pressure and reduced temperature, enabled shelf-life extension of raw milk for up to 5 weeks (the longest storage period studied). At -5°C/75 MPa, a volatiles profile and lipid oxidation levels similar to fresh raw milk were retained at 5 weeks, and there were no physiochemical or microbiological indications that substantially longer storage should not be achievable.

Comparing to available data on room-temperature hyperbaric storage at 75 MPa and 100 MPa [13], the combined high-pressure low-temperature conditions offered by isochoric freezing appear to reduce unfavorable changes in lipid oxidation and volatiles profile in raw milk, though direct experimental comparison would be needed to definitively conclude as much. Furthermore, hyperbaric storage requires active mechanical induction of pressure, whereas isochoric freezing produces passive self-pressurization driven only by temperature. As such, this preservation technique may prove more practical and adaptable to existing milk production and storage processes— low-temperature control infrastructure is already omnipresent within the dairy processing and distribution industry, and thus the only additional equipment necessary for this treatment would be isochoric storage containers.

As a whole, this study indicates that isochoric freezing has the potential to significantly increase the shelf life of milk by reducing microbial activity, whilst maintaining its nutritional and physiochemical content. The application of isochoric freezing, with its potential to adjust milk’s shelf life without significant deviation in quality, could offer a new means of reducing milk waste driven by temporal market fluctuations, and addressing future milk capacitance uncertainties driven by climate change. Future work might include isochoric treatment of non-raw milk products, examination of the absolute limits of milk shelf life under isochoric conditions, and study of the evolution of milk bacterial profile following prolonged isochoric storage.

## Acknowledgements

This work was supported by the USDA National Institute of Food and Agriculture, AFRI project Proposal #:2021-09570, Award #2022-67017-37098 “Novel Isochoric Processing for Sustainable, Safe and High-Quality Preservation of Fluid Foods”. A.M. also recognizes support from the National Science Foundation Graduate Research Fellowship Program under Grant No. DGE 2146752. Any opinions, findings, and conclusions or recommendations expressed in this material are those of the author(s) and do not necessarily reflect the views of the National Science Foundation.

## Data Availability

All data reported in this study is available from the authors upon reasonable request.

